# A predicted developmental and evolutionary morphospace for grapevine leaves

**DOI:** 10.1101/2022.01.29.478336

**Authors:** Daniel H. Chitwood, Joey Mullins

**Author notes:** Correspondence: Daniel H. Chitwood.

## Abstract

Using conventional statistical approaches there exist powerful methods to classify shapes. Embedded in morphospaces is information that allows us to visualize theoretical leaves. These unmeasured leaves are never considered nor how the negative morphospace can inform us about the forces responsible for shaping leaf morphology. Here, we model leaf shape using an allometric indicator of leaf size, the ratio of vein to blade areas. The borders of the observable morphospace are restricted by constraints and define an orthogonal grid of developmental and evolutionary effects which can predict the shapes of possible grapevine leaves. Leaves in the genus *Vitis* are found to fully occupy morphospace available to them. From this morphospace we predict the developmental and evolutionary shapes of grapevine leaves that are not only possible, but exist, and argue that rather than explaining leaf shape in terms of discrete nodes or species, that a continuous model is more appropriate.

## INTRODUCTION

Leaf shape across plants is diverse and spectacular, but it is not random. Development, evolution, and the environment sculpt leaf shape in specific ways (Chitwood and Sinha, 2016). Leaves allometrically expand, first shown by Stephen Hales through pin pricks on developing fig leaves that were displaced differentially along the length versus the width of the leaf (1727). The developmental programing of leaves changes from node-to-node resulting in changing leaf shapes. Goethe described this process as “metamorphosis” and in terms of the mutable, changing internal state of leaves (1817). Environment modulates leaf size and serrations, as observed by Bailey and Sinnott (1915) who used the distribution of entire leaves across latitudes to estimate the temperatures of paleoclimates. If we measure leaf shape across the seed plants, clear demarcations between phylogenetic groups are observed (Li et al., 2018). We have measured enough leaf shapes to know the borders and demarcations of what exists and the processes that shape leaves in specific ways.

The shapes of grapevine leaves have been measured under intense scrutiny and with purpose. Originally through morphometric techniques developed by Louis Ravaz (1902), the field of ampelography (“vine” + “process of measuring”) sought to discern, using leaves and other features of the vine, American *Vitis* species that were new to Europeans and would eventually be used as rootstocks against Phylloxera. Eventually the techniques would be famously applied to wine grape varieties by Pierre Galet (1979; 1985; 1988; 1990; 2000; Chitwood, 2020). Morphometric techniques have been used to genetically study the basis of leaf shape in grapevines (Chitwood et al., 2014; Demmings et al., 2019), how grapevine leaves develop (Chitwood et al., 2016a), the effects of environment (Chitwood et al., 2016b; Baumgartner et al., 2020), and to show that increases in vein length compensate for leaf area lost to lobing (Migicovsky et al., 2022a). Modeling has been used in several ways, including calculating average shapes of grapevine varieties while preserving features (Martínez et al., 1995; 1997a; 1997b; 1999), modeling development across grapevine shoots (Bryson et al., 2020), and using leaf allometry, specifically the ratio of vein to blade areas, as a proxy of leaf size and to measure the effects of year-to-year variation in leaf shape (Chitwood et al., 2021). For grapevines, as for many other types of leaves, we have extensively measured and modeled leaf shape, allowing us to discern genetic, developmental, and environmental effects with great power.

But what about leaves that are not available for us to measure? Using what we know about the underlying structure of leaf morphospaces across genotypic, developmental, and environmental effects, and making modeling assumptions about what is and is not possible, could we compare what we have measured and observed against the boundaries of what we know is possible?

Here, we measure the shapes of over 8900 grapevine leaves and model them against an allometric indicator of leaf size, vein-to-blade ratio, across *Vitis* species. The expansion of blade area at the expense of that for veins is found to be a principal determinant of the resulting morphospace, as much so as differences in leaf shape between species. These developmental and evolutionary forces that sculpt leaf shape are independent and lie orthogonal to each other. Using an inverse transform of the Principal Component Analysis (PCA) space, theoretical leaves missing from the data are reconstructed. We find that the borders of the grapevine leaf morphospace are sharply defined by developmental constraints of lobing and the ratio of vein-to-blade area and that leaves in the genus *Vitis* fully occupy the space available to them. Rather than discrete stages of development or species, for leaf shape, the morphospace is better described continuously as a grid defined by developmental and evolutionary effects from which any leaf shape in the genus *Vitis* can be predicted.

## MATERIALS AND METHODS

This work uses two sources of genetic material to sample grapevine leaf shape, referred to as “New York germplasm” and “California populations”. The first is the USDA germplasm repository in Geneva, NY which samples mostly North American *Vitis* species leaves (although not exclusively) as a developmental series, keeping track of the node the leaves arise from. These leaves tend to be more entire (again, not exclusively so). The second source of materials are segregating populations in California from E. & J. Gallo Winery (the exact identity of which is proprietary). The parentage of this material arises from *Vitis vinifera, V. mustangensis,* and *V. piasezkii* species and is more deeply lobed than the New York germplasm material (again, this is not always the case). Only mature, fully expanded leaves from the middle of the shoot were sampled from this population. This population was not sampled as a developmental series and the node the leaves arise from was not recorded. The New York germplasm allows models of leaf development to be estimated whereas the California populations sample additional leaf shapes throughout the genus *Vitis.* More specific information about each of these materials is given below.

## New York germplasm material

As described in Bryson et al., 2020 (and copied verbatim here for convenience), leaves were collected from 209 vines at the USDA germplasm repository vineyard in Geneva, New York, USA. Samples were taken from the same vines during the second week of June, annually, in 2013 and 2015–2017. The vines sampled represent 11 species *(Ampelopsis glandulosa* (Wall.) Momiy. var. *brevipedunculata* (Maxim.) Momiy., *V. acerifolia* Raf., *V. aestivalis* Michx., *V. amurensis* Rupr., *V. cinerea* (Engelm.) Millardet, *V. coignetiae* Pulliat ex Planch., *V. labrusca* L., *V. palmata* Vahl, *V. riparia* Michx., *V. rupestris* Scheele, and *V. vulpina* L.), four hybrids

(*V.* ×*andersonii* Rehder, *V.* ×*champinii* Planch., *V.* ×*doaniana* Munson ex Viala, and *V.* ×*novae-angliae* Fernald), and 13 *Vitis* vines, designated as *Vitis* spp., for which original species assignments from the germplasm collection are lacking. Starting at the shoot tip (with shoot order noted for each leaf), leaves greater than ~1 cm in length were collected in stacks and stored in a cooler in labeled plastic bags with ventilation holes. Within two days of collection, the leaves were arranged on a large-format Epson Workforce DS-50000 scanner (Tokyo, Japan) in the order they were collected, with a small number near each leaf indicating which node it came from and a ruler for scale within the image file. The image files were named with the vine identification number, followed by a sequential lowercase letter if multiple scans were needed. The original scans are available on Dryad (Chitwood et al., 2020).

## California populations material

As described in Migicovsky et al., 2022a (and copied verbatim here for convenience), leaves were sampled from seedlings of five biparental *Vitis* populations located in Madera County, California, USA. 500 seedlings were planted in the vineyard. 450 seedlings shared a seed parent, DVIT 2876. The remaining 50 seedlings had DVIT 2876 as a grandparent. DVIT 2876 ‘Olmo b55-19’ is a compound-leafed accession from the USDA-ARS National Clonal Germplasm repository, suspected to include *V. piasezkii* Maximowicz, as one of its parents (or grandparents). The populations were created to examine variation in leaf lobing. The vines were composed of 125 individuals from a DVIT 2876 x unnamed *V. vinifera* selection cross (Pop1), 100 individuals from a DVIT 2876 x a different unnamed *V. vinifera* selection cross (Pop2), 150 individuals from a DVIT 2876 x unnamed *Vitis* hybrid cross (Pop3), 75 individuals from a DVIT 2876 × a different unnamed *Vitis* hybrid cross (Pop4), and 50 individuals from a seedling (DVIT 2876 x unnamed *V. vinifera* selection) × DVIT 3374 (*V. mustangensis* Buckley) cross (Pop5). The vines sampled were planted in 2017. They were trained to a unilateral cordon and spur pruned. Leaf samples were collected on June 22 and July 12 2018, then again in 2019 on June 14, 19, and July 4. Across the sampling dates within a given year, a total of three mature, representative leaves were sampled from each of the vines and placed into labeled plastic bags. The plastic bags were stored in a cooler during collection and scanned, abaxial side down, later the same day using a flatbed scanner. Files were named using the accession identification number. The original scans are available on Dryad (Migicovsky et al., 2022b).

## Data Analysis

Twenty one landmarks (**Figure 1A**) were placed on one half of each leaf outlining the midvein, distal vein, proximal vein, and the most proximal branching vein of each of these major veins as well as distal and proximal lobe sinuses using ImageJ (Abràmoff et al., 2004). Two landmarks are placed at the base of each vein to measure the width. Landmarks were superimposed through scaling, translation, rotation, and reflection using Generalized Procrustes Analysis with the shapes (Dryden and Mardia, 2016) package in R.

**Figure 1:**
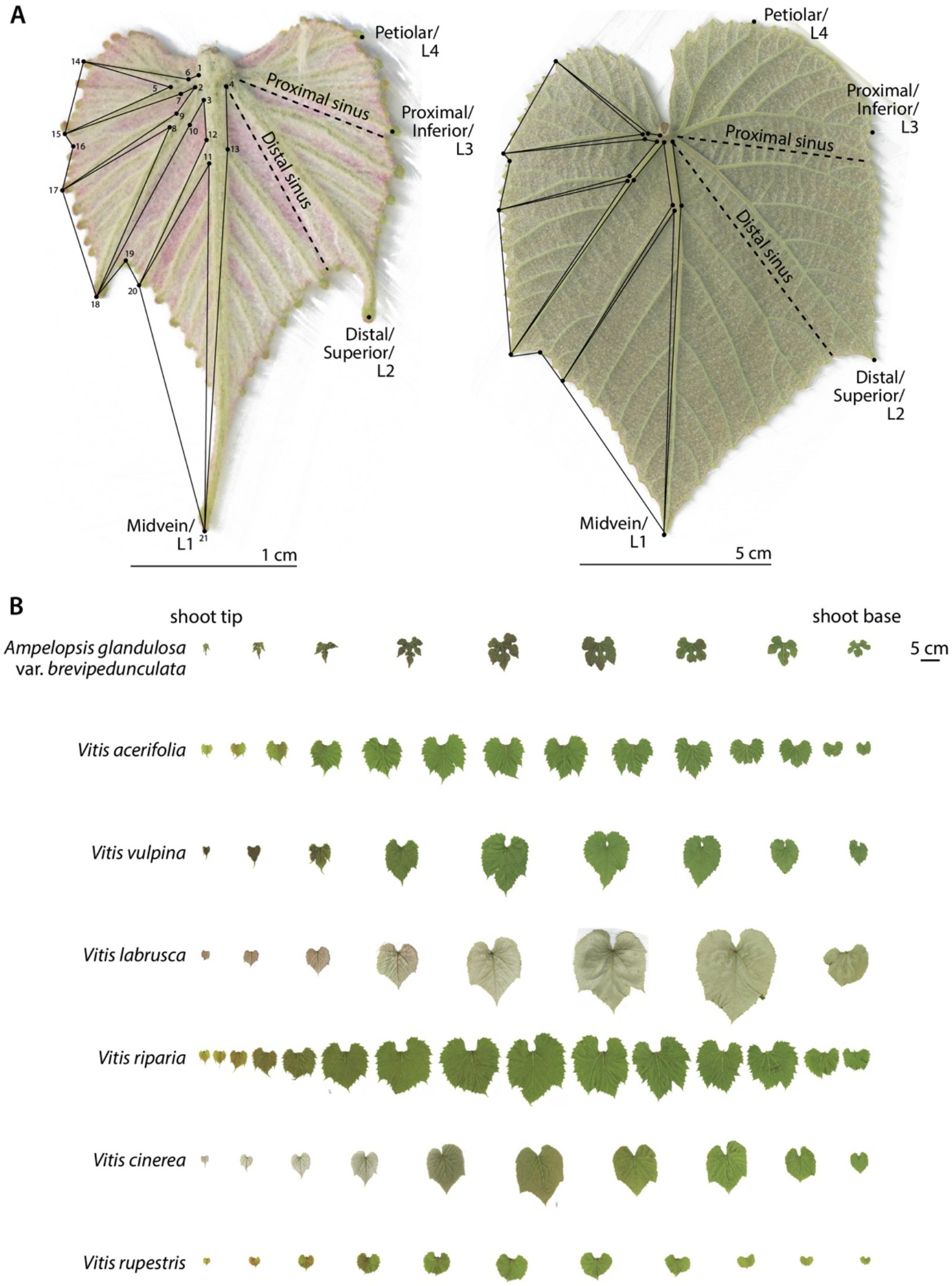
Grapevine leaf morphology. **A)** Counting from the shoot tip, *Vitis cinerea* leaves from node positions 1 (left) and 5 (right), each with respective scale bar, are expanded in detail from the same leaves shown in the panel below. The 21 landmarks used in this study are indicated, as well as ampelographic nomenclature naming morphological features. Note that in the younger leaf that vasculature takes up relatively more area than in the mature leaf. **B)** For seven different grapevine species analyzed in this study, leaves from the shoot tip to the shoot base are shown with scale bar. Leaf area increases from the shoot tip to the middle of the shoot due to leaf expansion, whereas increases in leaf size from the shoot base to the middle of the shoot in mature leaves are due to heteroblasty.

Data was analyzed using Python and Jupyter notebooks (Kluyver et al., 2016). Code to reproduce the analysis in this manuscript can be found at the *Github* repository DanChitwood/grapevine_morphospace: https://github.com/DanChitwood/grapevine_morphospace). The Jupyter notebook (grapevine_morphospace.ipynb) comments on the code and also contains a narrative to guide the reader through the analysis. Calculation of distal lobing is according to Galet (1979), as the ratio of the distance of the distal sinus to the petiolar junction divided by the distance of the distal lobe tip to the petiolar junction, such that the distal lobing value of a completely dissected leaf is 0 and the value of a completely entire leaf is 1.

Calculation of the natural log of the ratio of vein to blade area, ln(*vein to blade ratio*), is as described in Chitwood et al. (2021) using the shoelace algorithm, also known as Gauss’ area formula, to calculate polygon areas as originally described by Meister (1769), where *n* is the number of polygon vertices defined by *x* and *y* coordinates:

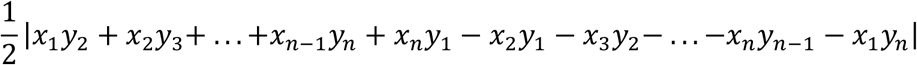

Principal Component Analysis (PCA) (and calculation of its inverse) was performed using the scikit learn decomposition PCA module (Pedregosa et al., 2011). Modeling of ln(*vein to blade ratio*), ln(*leaf area*), and landmarks as polynomial functions of each other and shoot position was performed using the np.polyfit and np.poly1d functions from NumPy (Oliphant, 2006). The curve_fit function from SciPy (Virtanen et al., 2020) was used to fit a reciprocal function of ln(*leaf area*) across the shoot. Pandas (McKinney, 2010) and Matplotlib (Hunter, 2007) were used for data analysis and visualization.

## RESULTS

### Developmental models of leaf expansion

Previously, we modeled leaf shape continuously across grapevine shoots as a polynomial function of each Procrustes landmark coordinate value as a function of normalized node position. Normalized node position is the node number counting from the shoot tip divided the total number of leaves in a shoot, such that node number is converted to a 0 to 1 scale, from tip to base (Bryson et al., 2020). We also previously described the natural log of the ratio of vein to blade area, ln(*vein to blade ratio*), which is more sensitive to leaf area than size itself due to the exponential increases in blade relative to vein area during development (Chitwood et al., 2021). Before we explore the limits of the grapevine leaf morphospace, we must first model shape across development to understand how continuous developmental trajectories change between species during evolution. But it is important to first understand two developmental processes that affect leaf size and shape across grapevine shoots. At the shoot tip and base leaves are smaller (and accordingly ln(*vein to blade ratio*) is higher) than the middle of the shoot where leaves are larger (and ln(*vein to blade ratio*) lower) (**Figure 1B**). At the shoot tip leaves are young and at the shoot base they are mature. The increases in leaf area (and decreases in ln(*vein to blade ratio*)) from the shoot tip to the middle of the shoot are mostly due to the expansion of young leaves as they mature. However, the increases in leaf area (and decreases in ln(*vein to blade ratio*)) from the shoot base to the middle of the shoot occur in mature leaves that have already expanded. The size and shape differences between mature leaves at the shoot base are due to heteroblasty, node-to-node differences in leaf morphology that result from the temporal development of the shoot apical meristem, and not from leaf expansion.

Below, we create models of leaf development to focus on allometric changes due to leaf expansion and its relationship to the grapevine leaf morphospace. To do so requires us to separate these confounding effects on leaf shape and size across the grapevine shoot to the best of our ability.

We plotted ln(*vein to blade ratio*) versus normalized node position (**Figure 2A**), which can be modeled as a second-degree polynomial. ln(*vein to blade ratio*) is highest at the shoot tip and reaches its minimum in the middle of the shoot. As expected, In(*leaf area*) versus relative node position correspondingly increases in the middle of the shoot compared to the shoot tip and base (**Figure 2B**). A curiosity that is perhaps coincidental, we note that the corresponding normalized node position to the minimum ln(*vein to blade ratio*) and maximum ln(*leaf area*) values are close to the inverse of the golden ratio (**Figure 2A–B**). Although this may arise as a developmental phenomenon, it could also be spurious and warrants further investigation.

**Figure 2:**
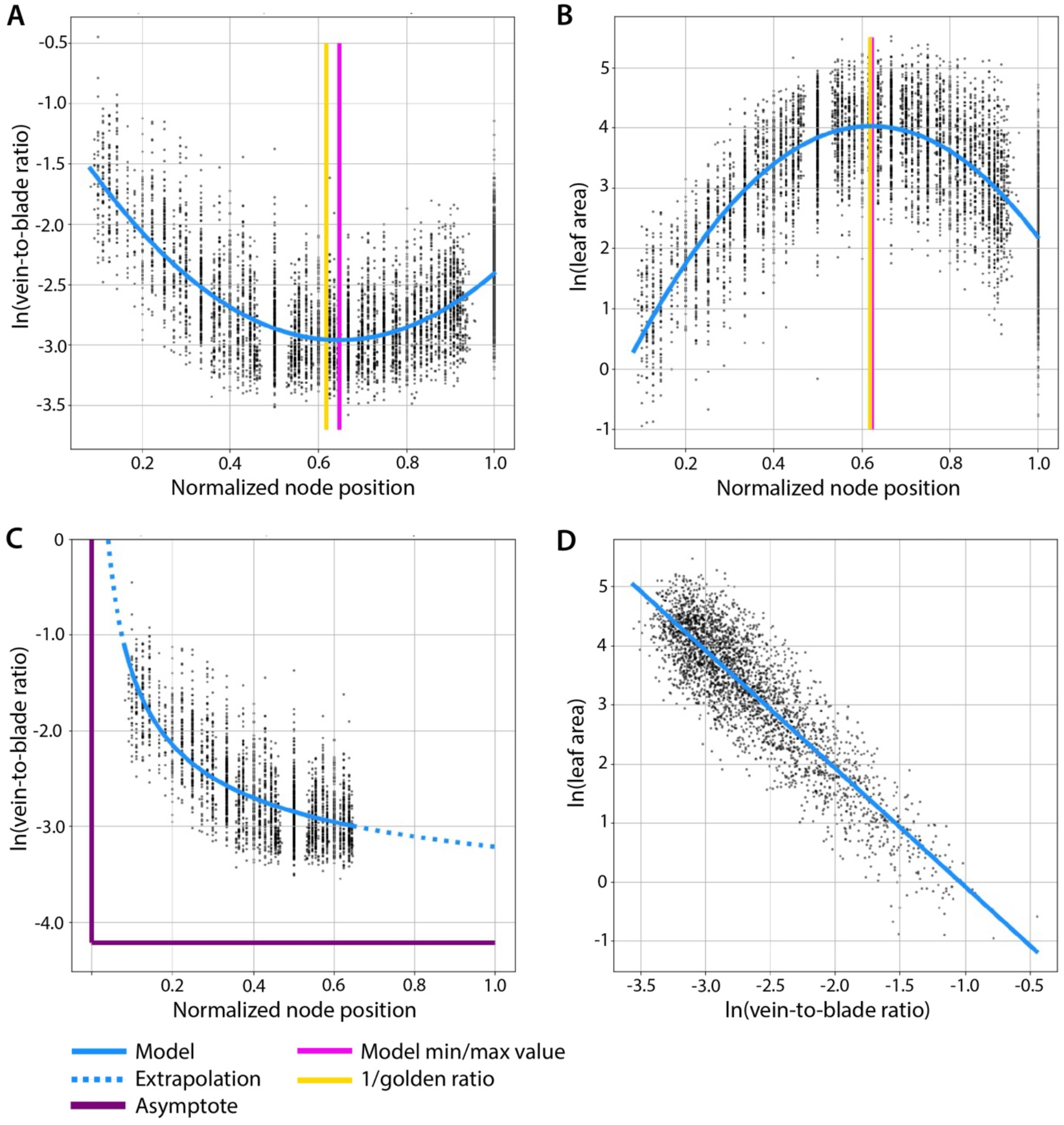
Modeling ln(vein *to blade ratio*) and *ln*(*leaf area*) as a function of normalized node position. **A)** The natural log of the ratio of vein-to-blade area, *ln(vein to blade ratio),* and **B)** the natural log of leaf area, ln(*leaf area*), are modeled as 2nd degree polynomials of normalized node position (where 0 is the shoot tip and 1 is the shoot base). The normalized node position values corresponding to the minimum ln(*vein to blade ratio*) and maximum ln(*leaf area*) values are indicated by a magenta vertical line and the inverse of the golden ratio is indicated by a gold vertical line. **C)** In order to model developmental changes due to leaf expansion separate from heteroblastic effects, leaves from the shoot tip to the normalized node position value corresponding to the ln(*vein to blade ratio*) minimum were isolated and modeled as a reciprocal function of normalized node position. Extrapolated values are shown in dashed line and function asymptotes in purple. **D)** A linear model of ln(*leaf area*) as a function of ln(*vein to blade ratio*).

From previous work we know that allometric changes during grapevine leaf expansion dominate the morphospace (Chitwood et al., 2016a; Chitwood et al., 2016b; Bryson et al., 2020). We therefore took leaves from the shoot tip to the normalized node position value corresponding to the minimum ln(*vein to blade ratio*) value across the shoot (**Figure 2A**) to model shape changes associated with leaf expansion. Assuming that ln(*vein to blade ratio*) approaches ∞ as a normalized node position value of 0 is approached (leaf initiation, where vein area would dominate) and that another asymptote is approached as leaves mature (where blade area dominates) a reciprocal function was fit to the data (**Figure 2C**). Using the model, the context of the collected data compared to extrapolated leaf shapes that remain unsampled (for example, young leaf primordia or leaves that continue to mature incrementally past the leaves collected in this study) can be understood. From these expanding leaves a linear model of ln(*leaf area*) as a function of ln(*vein to blade ratio*) can be fit (**Figure 2D**). From this model, using a scaleless measure of leaf shape alone, leaf size can be predicted. Importantly, for the expanding leaves selected for modeling above, their ln(*vein to blade ratio*) values are always decreasing, and their leaf area values are always increasing moving away from the shoot tip, separating and unconfounding these effects from those of heteroblasty (**Figure 1B**).

By modeling Procrustes-adjusted coordinate values as a polynomial function of ln(*vein to blade ratio*), we can visualize and compare the developmental trajectories of different grapevine species (**Figure 3**). Theoretical leaves for the six most represented *Vitis* species and *Ampelopsis glandulosa* var. *brevipedunculata* across ten equally spaced ln(*vein to blade ratio*) values from the maximum to minimum (inclusive), show the shape changes associated with leaf expansion and evolutionary differences between species. Leaf expansion is mostly achieved through increases in blade area relative to vein, as well as other changes, such as a wider leaf. These developmental changes in shape are conserved and distinct from species differences, which affect a different set of shape features, especially the depth of the distal lobe. These shape changes are allometric and occur concomitantly with exponential decreases in leaf size. The developmental models of leaf expansion described above will be projected onto the morphospace described below to anchor and contextualize the space and to quantify and compare evolutionary versus developmental sources of shape variance across grapevine leaves.

**Figure 3:**
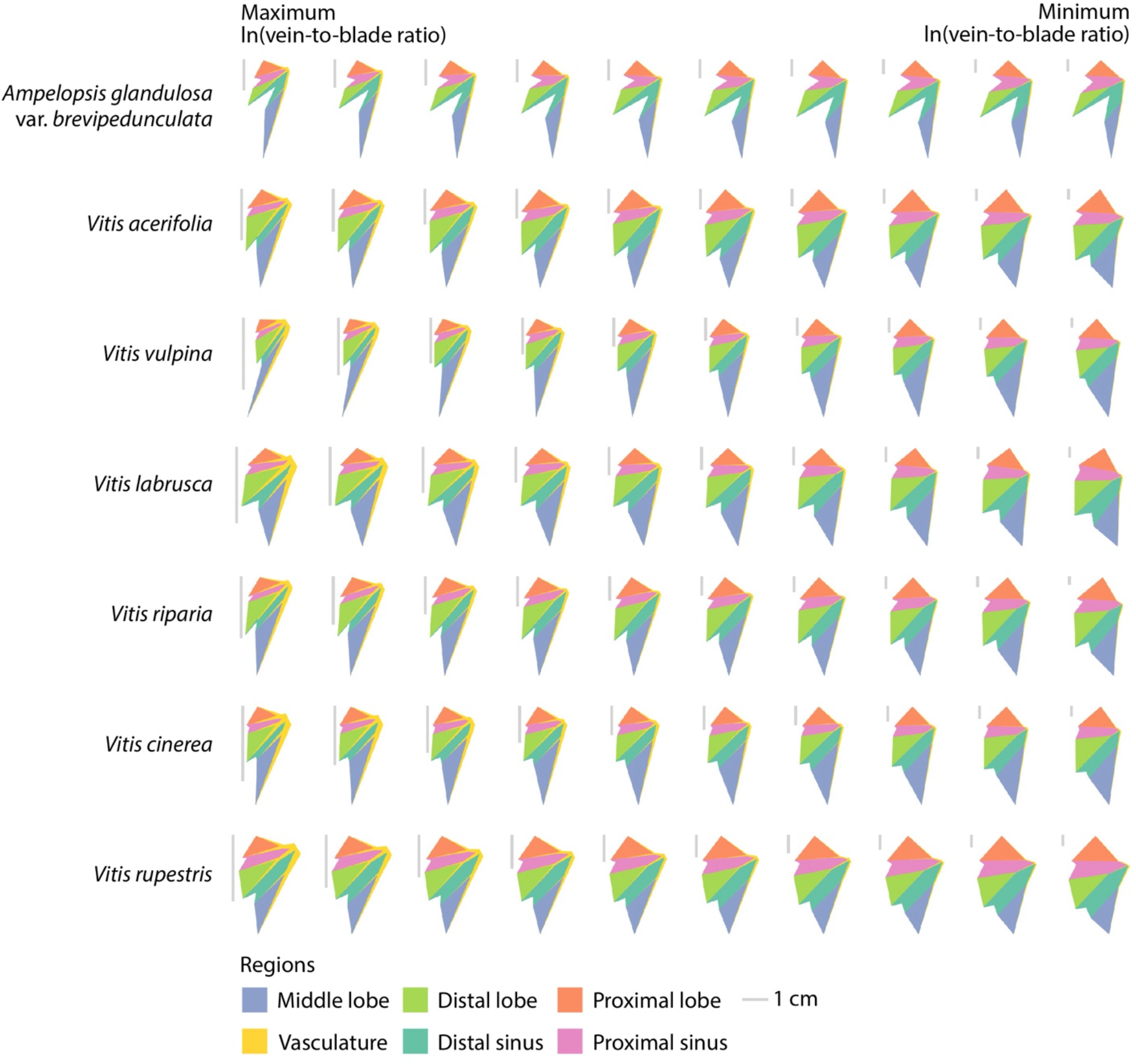
Developmental models of leaf shape. Fitting each coordinate value of 21 landmarks as a second-degree polynomial of ln(*vein to blade ratio*), continuous models of expanding leaves for the seven species shown were created. Inclusive of the maximum and minimum ln(*vein to blade ratio*) values for each species, corresponding to young and mature leaves, respectively, leaves corresponding to ten equally spaced time points were reconstructed. Estimated leaf areas were estimated from ln(*vein to blade ratio*) values and 1 cm scale bars for each leaf are shown. Leaf areas are indicated by color.

### Morphospace

The developmental models of leaf expansion described above are from a dataset, the “New York germplasm”, where leaves were sampled from shoots and their node position was recorded. These leaves, from the USDA germplasm repository in Geneva, NY sample mostly (although not exclusively) North American *Vitis* species that tend to have more entire leaves (although there are highly dissected leaf samples in the dataset). Largely missing is shape variation from *V. vinifera* and other highly dissected species. To supplement the New York germplasm leaves, we added leaves from segregating populations designed to sample highly lobed genetic material, derived from *V. vinifera, V. mustangensis,* and *V. piasezkii,* called the “California populations”. All leaves from the California populations are mature, creating an opportunity to predict and extrapolate the development of these leaves from the New York germplasm. Although not representing the entirety of mature leaf shape variation within *Vitis,* the two datasets together comprehensively sample it.

To visualize the relationship of New York germplasm to California populations datasets, and how developmental versus evolutionary sources of leaf shape variation compare, we performed a Principal Component Analysis (PCA). PCA decomposes multivariate data, in essence rotating and projecting it onto orthogonal axes (principal components) that more efficiently explain variation in the data than the original measurements (in this case, Procrustes-adjusted coordinate values). The inverse of this transformation can be used to reverse calculate original data, which we will later use to visualize theoretical leaves in the morphospace. PC1 and PC2 explain 39.7% and 17.6% of the variance in the data, respectively (~57.3% of the total variance). Within this space, the NY germplasm and CA population data are roughly orthogonal (perpendicular) to each other (**Figure 4**). One interpretation is that the more entire leaves of the NY germplasm data run along a developmental continuum, whereas the California populations data only represents mature leaves but falls on a separate axis representing leaves that are more dissected. The empty space not covered within the ranges of the two datasets would be predicted to be the missing developmental variation from the deeply lobed leaves in the California populations data. Two pieces of evidence support the above interpretation. First, if developmental models of leaf expansion are projected onto the morphospace, they are collinear with the distribution of the New York germplasm data, consistent with this axis of the data representing developmental variation. Second, if ln(*vein to blade ratio*) values for theoretical leaves calculated from the inverse transform of the morphospace are projected back onto it (**Figure 4A**) they too are collinear with the NY germplasm data. Similarly distal lobing, which varies across species (**Figures 1 and 3**), can also be calculated and projected back onto the morphospace (**Figure 4B**). Distal lobing runs at roughly right angles to ln(*vein to blade ratio*) values and the CA populations data is collinear with it. The CA populations data intersects with the NY germplasm data in a location defined by low ln(*vein to blade ratio*) values, consistent with these being mature leaves.

**Figure 4:**
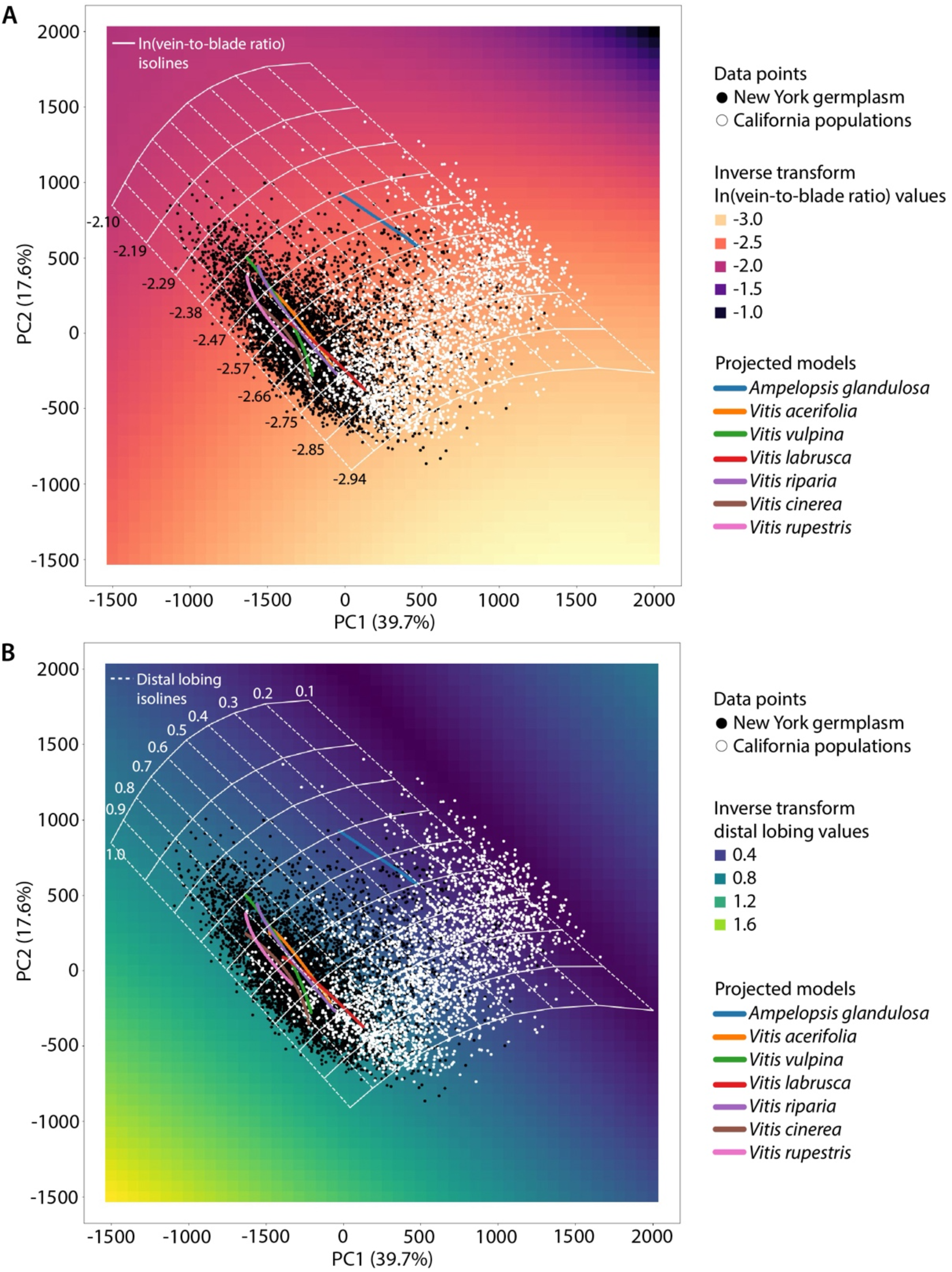
Morphospace. A morphospace calculated from a Principal Component Analysis (PCA) of all leaves from the New York germplasm (black) and California populations (white). **A)** ln(*vein to blade ratio*) values and **B)** distal lobing values were calculated from reconstructed leaves throughout the morphospace using its inverse transform and colored by magma and virdis color schemes, respectively, as indicated. To orient and contextualize the space, developmental models for seven grapevine species were projected into the space, as indicated by colored lines. Isolines for **A)** ln(*vein to blade ratio*) values (solid lines) and **B)** distal lobing values (dashed lines) are shown and their values provided in the respective plots.

If developmental variation (indicated by ln(*vein to blade ratio*) values, **Figure 4A**) and evolutionary variation between species (indicated by distal lobing values, **Figure 4B**) are roughly orthogonal to each other, then even though unsampled, the shapes of developing leaves that are highly dissected that are missing from the CA populations data could be predicted. The ability to make this prediction rests on the assumption that highly dissected leaves would follow a developmental trajectory similar to more entire species. Evidence that this is the case is observed for the developmental model of *Ampelopsis glandulosa* var. *brevipedunculata* (**Figure 4**), which is collinear like the other models with ln(*vein to blade ratio*) values and occupies a space with low distal lobing values, consistent with its deeply lobed morphology.

Beyond stages of leaf development and different species, the morphospace of grapevine leaves can be described more quantitatively and comprehensively using ln(*vein to blade ratio*) and distal lobing values that define it continuously. Isolines that fall along the same ln(*vein to blade ratio*) and distal lobing values can be calculated so that they extend to the borders of observable morphospace and sample, in a grid-like fashion, the space inside. These isolines also sample inferred leaf shapes not represented in the sampled data, including the missing developmental series from the CA populations data and leaf primordia younger than those sampled. Theoretical, reconstructed leaves at the intersection of ln(*vein to blade ratio*) and distal lobing isolines, that sample the limits of the observable morphospace, exhibit the distinct changes in shape associated with development and evolution (**Figure 5**). Across developmental series regardless of how deeply lobed leaves are, ln(*vein to blade ratio*) decreases and leaves become wider as they expand and increase in size. Similarly, as ln(*vein to blade ratio*) isolines traverse orthogonally to distal lobing isolines, the depth of the distal lobe is preserved regardless of developmental stage and comprises evolutionary differences in grapevine leaf shape that are independent of development.

**Figure 5:**
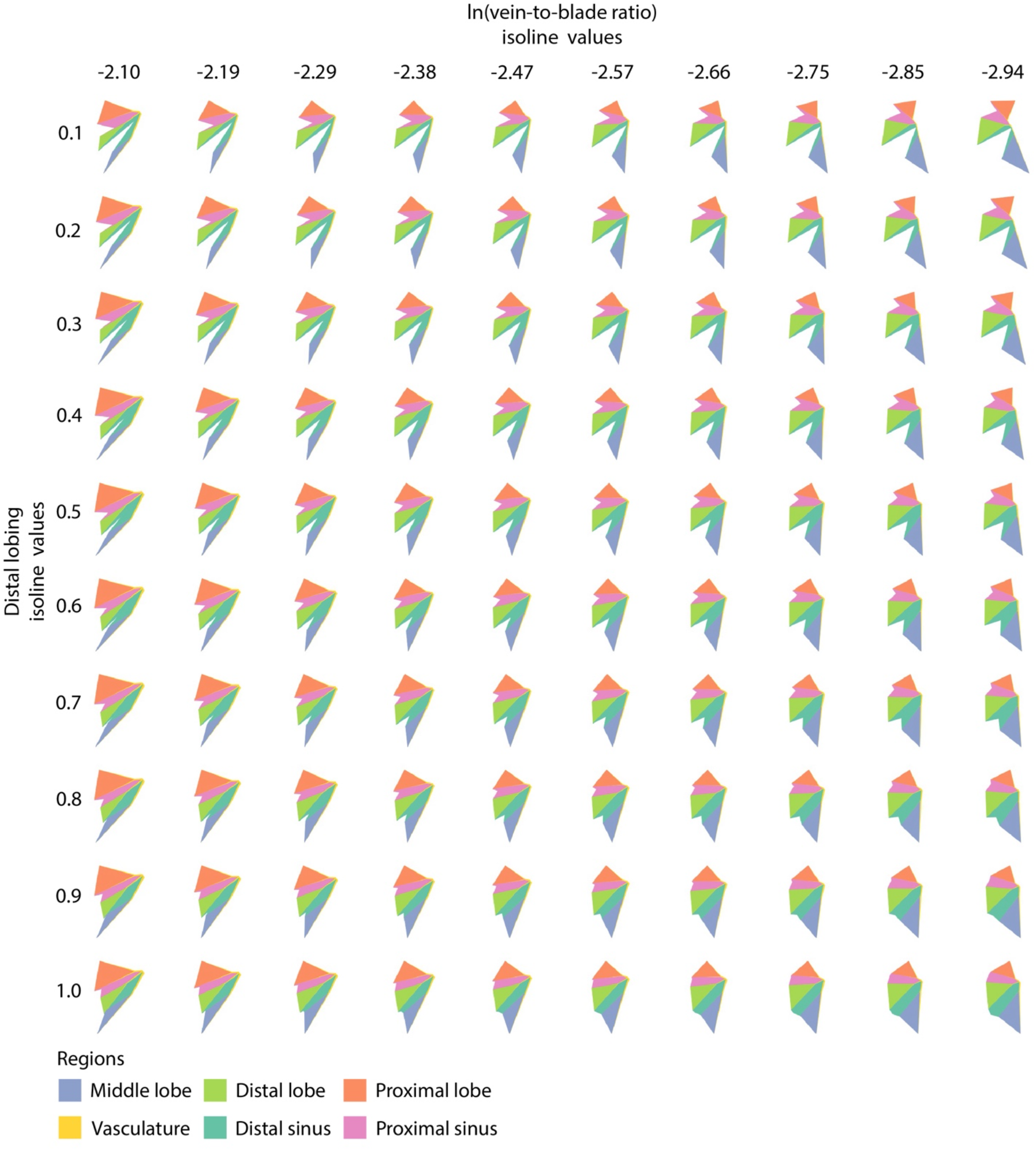
Theoretical leaves. 100 theoretical leaves reconstructed from the intersection of ten, equally spaced ln(*vein to blade ratio*) and distal lobing isolines, corresponding to orthogonal developmental and evolutionary changes, respectively, across grapevine leaf morphospace. ln(*vein to blade ratio*) and distal lobing values are shown and leaf areas indicated by color.

## DISCUSSION

PC1 and PC2 together explain round 57.3% of the variance in the data, but they represent the first two major, orthogonal sources of variance and as described (**Figure 4**) highlight natural axes in the data that delimit developmental and evolutionary boundaries that constrain observable grapevine leaf shapes. ln(*vein to blade ratio*) and distal lobing are only indicators of multivariate signatures of leaf development and evolution, respectively, that lie orthogonal to each other and define a grid in which grapevine leaves fully occupy to its limits. One set of boundaries is indicated by distal lobing values (dashed isolines in **Figure 4B**), defined by leaves with values approaching zero and completely dissected (like *A. glandulosa* var. *brevipedunculata* or *V. piasezkii)* or nearly equal to one and lacking any significant lobing (like *V. rupestris*). The other set of boundaries is indicated by ln(*vein to blade ratio*) values (solid isolines in **Figure 4A**) that asymptotically define developmental constraints. Higher ln(*vein to blade ratio*) values are associated with young, expanding leaves in which vein area initially dominates the leaf until the blade exponentially expands. The developmental models presented in this analysis work from the assumption that young leaf primordia approach an asymptote consisting entirely of vein area at initiation (**Figure 2C**). In leaves that are nearly fully expanded the opposite is true, and they are defined by lower ln(*vein to blade ratio*), in which a small amount of vein area remains, but that blade will always allometrically expand at a faster rate than vein and approach an asymptote in which vein area is vanishingly small (**Figure 2C**).

The morphospace is unexpectedly simple, providing a predictive framework and empirical insight into theoretical biological concepts. While the New York germplasm and California populations data sample most shape variation in *Vitis,* the developmental information for highly dissected species was missing. Because developmental and evolutionary axes are nearly orthogonal to each other and describe additive signatures of leaf morphology, where developmental progressions in leaf shape are conserved across species and variation defining differences between species is maintained throughout their development, to extrapolate the leaf shapes missing in this space was straightforward (**Figure 5**). In theory we talk about evolutionary and developmental forces describing the organismal form, but definition is lacking: to what degree do they act separately or are confounded together, do they act additively or do interaction effects predominate? In the case of grapevine leaves, development and evolution are orthogonal and acting independently of each other to such an extent that rather than describe leaf shape as arising from discrete nodes or species, a continuous model defined by indicators like ln(*vein to blade ratio*) and distal lobing is more efficient (**Figure 4**). It is also an open question to what degree developmental constraint and selection would limit the full manifestation of phenotype across a morphospace. For the example of grapevine leaf shape, the boundaries of the morphospace are well defined by developmental constraint and it appears that development and evolution have fully sampled the space, up to the borders (**Figure 4**).

Although reconstructing leaves from a PCA morphospace is routine statistically, this work focuses on interpretation and how we can use morphometrics to see shape and natural phenomena through different lenses. Embedded in the morphology of morphospaces we measure are the constraints by which development and evolution are modulating natural forms. Measured in sufficient quantities and making reasonable assumptions about the limits of our models, we can begin to deduce and quantify constraint, and predict the extent of what is phenotypically possible.

## DATA AND CODING AVAILABILITY STATEMENT

The code to reproduce this analysis can be found at the *Github* repository DanChitwood/grapevine_morphospace: https://github.com/DanChitwood/grapevine morphospace. The original leaf scans used to produce the landmarks are archived on Dryad (Chitwood et al., 2020; Migicovsky et al, 2022b).

## AUTHOR CONTRIBUTIONS

JM developed methods and tools, performed data analysis, and reviewed the manuscript. DHC conceived of the project, performed data analysis, and wrote the manuscript.

